# Non-hypermutator cancers access driver mutations through reversals in germline mutational bias

**DOI:** 10.1101/2024.04.30.591900

**Authors:** Marwa Z. Tuffaha, David Castellano, Claudia Serrano Colome, Ryan N. Gutenkunst, Lindi M. Wahl

## Abstract

Cancer is an evolutionary disease driven by mutations in asexually-reproducing somatic cells. In asexual microbes, bias reversals in the mutation spectrum can speed adaptation by increasing access to previously undersampled beneficial mutations. By analyzing tumors from 20 tissues, along with normal tissue and the germline, we demonstrate this effect in cancer. Non-hypermutated tumors reverse the germline mutation bias and have consistent spectra across tissues. These spectra changes carry the signature of hypoxia, and they facilitate positive selection in cancer genes. Hypermutated and non-hypermutated tumors thus acquire driver mutations differently: hypermutated tumors by higher mutation rates and non-hypermutated tumors by changing the mutation spectrum to reverse the germline mutation bias.

## Introduction

Cancer is an evolutionary disease arising from DNA mutations that allow cells to proliferate abnormally and invade other tissues (*1*). During the asexual reproduction of normal somatic cells, *de novo* mutations accumulate through time, which increases cancer risk with age (*2*). Cancer develops when mutations in specific genes or combinations of genes (so-called “cancer genes”) impair normal cell function by, for example, disabling the cell cycle checkpoints or activating signals which drive excessive cell division (*3*).

For both evolution within a tumor and evolution within a population of asexual organisms, the mutation rate is a key factor. In microbial evolution, lineages with an increased mutation rate (mutators) frequently emerge due to their improved access to rare beneficial mutations (*4*). Along with increases in mutation rate, changes in mutation spectra also affect mutational supply (*5–7*). We previously demonstrated that the interaction between mutation rate and spectrum powerfully influences the evolutionary trajectory of asexual populations (*8*). Simply put, previously undersampled mutations may be reached by either making more mutations (mutation rate elevation, fig. 1A), or by making different mutations (mutation spectrum change, fig. 1B). If mutations have historically occurred with some bias (undersampling some classes of mutations while oversampling others), reversing this bias affords access to mutations that were previously unlikely to have occurred, such as undersampled beneficial mutations (*7*).

**Fig. 1.**
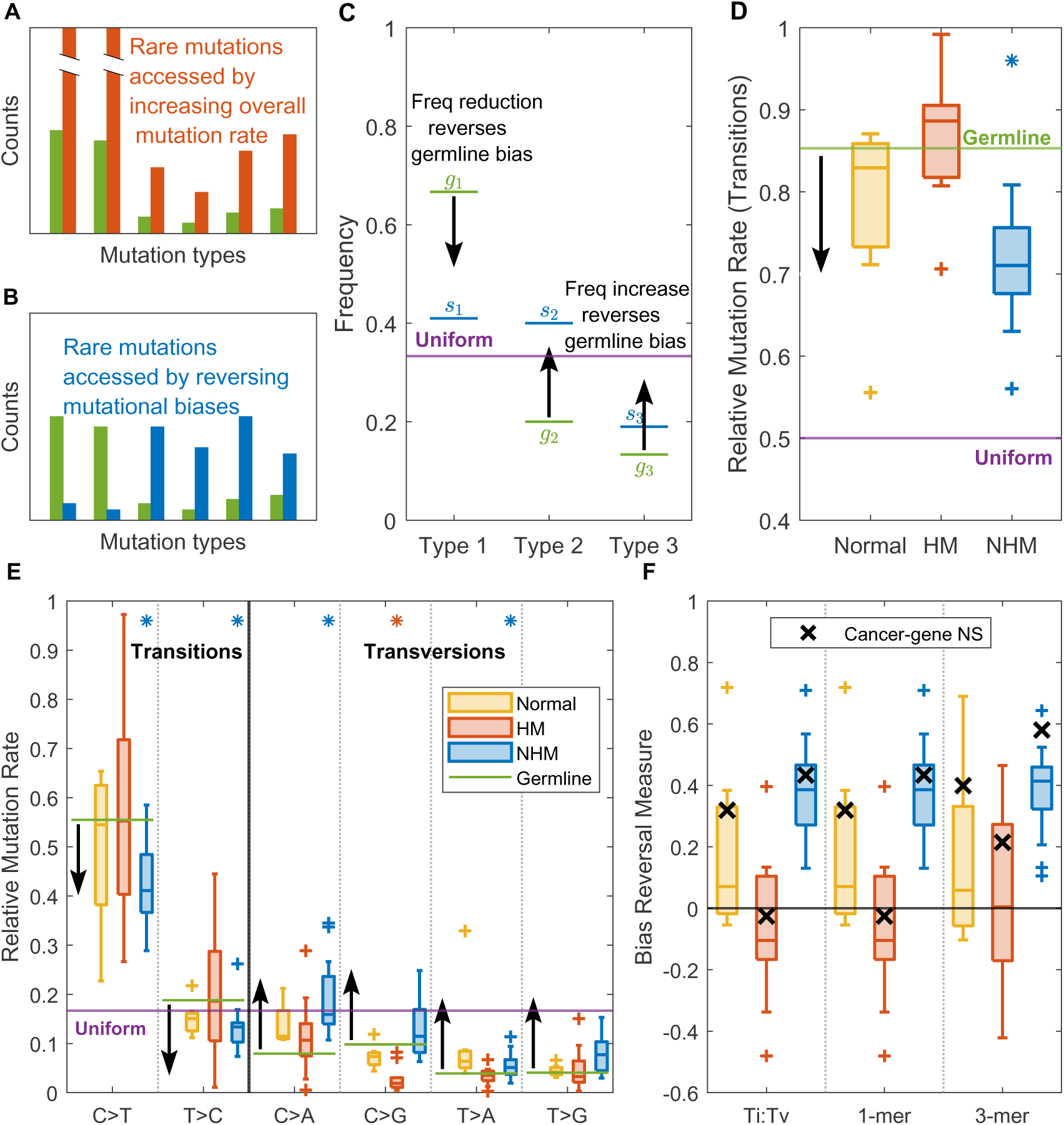
(A,B) Schematic: a germline spectrum (green) under-represents some classes of mutations. Access to these classes increases by either an overall mutation rate increase (A) or reversing the bias (B). (C) Example of a spectrum with 3 types of mutations having the same uniform frequency 1/3. Type 1 is over-represented in the germline (*g*_1_, green) while the other two are under-represented. Arrows show the direction that reverses the bias for each mutational type. An example of a bias-reversing spectrum is shown in blue (*s_i_*). (D) Distributions of the RMR for transitions in normal tissues (yellow), hypermutator tumors (HM, red) and non-hypermutator tumors (NHM, blue) in different tissues, compared to the uniform (purple) and germline (green) levels. (E) The corresponding RMRs for each 1-mer mutation type. Stars indicate distribution means that are significantly different from the germline. (F) Bias reversals (*y*-axis) for passenger gene spectra in different tissues (boxplots) are significantly higher in NHM than in HM or normal tissues at all three levels of analysis (*x*-axis). The bias reversal measure for non-synonymous mutations in cancer genes is also plotted (’x’). The germline results are shown as solid lines since the standard errors are smaller than the line thickness. Horizontal bars within boxplots indicate medians; whiskers indicate the 95% confidence interval (CI); ’+’ signs represent outliers.

Cancer is a set of diseases that have similar hallmarks (*9*), but there are many differences between cancer types, including different mutational processes and rates (*10*). Tumors vary in mutational burden both within the same tissue (across patients) and between different tissues (*11*). Hypermutation is usually caused by DNA mismatch repair defects, whether inherited or acquired from somatic mutations, which is common in gastrointestinal cancers (*12*). Exposure to UV light also elevates mutation rates in skin cancers (*13*). Thus, hypermutation is sometimes defined based on these etiologies rather than the exact number of mutations (*10*). There is debate over whether a single universal threshold can be established to define “hypermutation” for all cancer types and sequencing techniques, but it is generally considered to fall within the range of 10 to 20 mutations per megabase (mut/Mb) (*14–16*).

Along with these changes in mutation rate, biases in mutational spectra are widely observed in somatic mutations, where different tissues, individuals, and exposures show different tendencies for some types of mutations rather than the others to occur (*17*). Distinct patterns of mutations, called “mutational signatures”, arise in the DNA of normal (*18–20*) and cancer (*10, 21*) cells due to various biological processes, environmental exposures, or intrinsic factors. Whether and to what extent the combined effects of these diverse mutational processes exhibit mutational biases, and how these biases compare with the mutation bias of the germline has not yet been determined.

We investigate the interaction between mutation rate and mutation spectrum changes in cancer. In particular, we compare the mutation spectrum of hypermutator (HM) and non-hypermutator (NHM) cancers, demonstrating a highly conserved spectrum across NHM samples that consistently reverses the mutation bias of the germline. Cancer driver mutations thus occur through distinct mechanisms in HM and NHM cancers. In HM cancers the supply of cancer driver mutations is increased through higher mutation rates, while in NHM cancers, the reversal in mutation spectrum allows driver mutations. Our results emphasize the role of both the mutation rate and spectrum as critical in modeling genetic evolution in cancers and open new perspectives into the mechanisms that drive the development and progression of different cancers.

## Materials and Methods

### Datasets

We use three sets of data in our analysis, described as follows.

1. *PCAWG*: The Pan-Cancer Analysis of Whole Genomes (PCAWG) study (https:// dcc.icgc.org/pcawg) is an international collaboration which provides whole genome mutation data from cancer tissues from over 2,700 donors. Overall, the dataset has 55,657,793 mutations including 421,336 coding mutations, in 20 different primary tissues. Each sample in the PCAWG dataset was classified as a hypermutator cancer (HM) or a non-hypermutator cancer (NHM) based on the classification provided by (*10*). We note that the hypermutated set consists mainly of skin tumours and samples with mismatch repair or putative polymerase epsilon defects. The distribution of coding mutations from hypermutated and non-hypermutated samples in the different tissues is shown in fig. S1, while the distribution of whole-genome mutations as well as the number of donors in the dataset are clarified in figures S2 and S3. Effects and contexts of mutations were determined by mapping to the human genome hg19.
2. *Normal Tissue Mutations*: We collected whole-genome and coding mutations from 19 and 9 normal tissues as provided in the supplementary material of the work of (*18*) and (*19*), respectively.
3. *Germline de novo Mutations*: We consider the seven datasets used by (*22*), which consist of 679,547 germline single-base substitutions from family-based whole genome datasets from multiple centers in Europe, Asia and North America (see (*22*) for details). These include 10,750 coding mutations.

### Spectrum Calculation

We consider two types of spectra: absolute count spectra, and mutation rate spectra. The methods we use to calculate these spectra can be generalized to any spectrum with *N* mutational categories, and thus we explain for a general spectrum. However, we only calculate spectra at three different levels of detail: (1) the transition:transversion spectrum represented by a single measure, the transition frequency; (2) the 1-mer spectrum that includes the frequency of each 1-mer mutation type, and (3) the more detailed 3-mer spectrum, in which the mutation rate of each nucleotide depends on the identity of adjacent nucleotides.

Since a C*>*A mutation, for instance, on one strand of the DNA corresponds to a G*>*T mutation in the same position on the other strand, these two mutations are equivalent and we count them in a single mutational category; this category will be denoted C*>*A following the convention of taking the pyrimidine as the reference base. A 1-mer spectrum therefore consists of 6 categories of mutation, and the same idea leads to 96 categories in a 3-mer spectrum.

Assume *M* mutations are observed in a (sub)dataset, out of which *m_i_* are from a given mutational category *i*, so that its absolute frequency is *m_i_/M* . The vector of these absolute frequencies for all mutational types is what we call the *absolute count spectrum*.

When comparing spectra, it is sometimes useful to correct these absolute frequencies by the occurrence opportunities for each mutation type in the underlying genome. For example, a very high frequency of C*>*A mutations could be observed in a genome with high GC content, even if the underlying mutation rate is not elevated. Thus, to isolate changes in mutation rate, as opposed to genome content, we normalize the observed frequencies by mutational opportunities in the human genome and call such a spectrum the *mutation rate spectrum*. If mutational category *i* has *n_i_* opportunities to occur in the genome, then we define its *genomic mutation rate* as *m_i_/n_i_*. For 1-mer and 3-mer mutations, the factor *n_i_* is simply given by the genome content; for transitions and transversions, however, each base in the genome provides two opportunities for a transversion to occur, and one opportunity for a transition regardless of genome content. Normalizing the vector of these genomic mutation rates to add to one gives the mutation rate spectrum, i.e., the mutational frequencies if all categories had the same opportunity to occur, which will be referred to as the *relative mutation rates* as opposed to the absolute frequencies in the absolute count spectrum. We emphasize that both the absolute count spectrum and mutation rate spectrum are useful in different contexts, depending on whether we wish to compare the number of mutations (of each type) that actually occurred, or the underlying mutation rate of each type.

Genes that are recurrently mutated across cancer patients are identified as putative cancer driver genes. Signals of positive selection have been previously detected in a few hundred of such genes (*23*) using DNA sequencing data from tumor tissues. These signals measure an excess of amino-acid changing (non-synonymous) mutations compared to a null model. Because mutation hotspots could also be recurrently mutated across patients, several sophisticated methods that account for the peculiarities of somatic mutation spectra have been proposed (*24–26*). When analyzing the spectra of the coding genome, we exclude mutations that occur in cancer genes identified by the Cancer Gene Census list (*23*). Restricting our analysis to mutations in non-cancer “passenger” genes eliminates the possibility that mutations driven by positive selection in cancer genes might distort the spectrum.

When comparing spectra across tissues or cancer types, we directly compare the corresponding Transition:Transversion (Ti:Tv) and 1-mer frequencies. To compare 3-mer spectra, we use correlation coefficients. Since both cosine similarity and Pearson correlation yield results dominated by CpG transitions (3-mers in which C*>*T) due to their high frequencies, we use Spearman rank correlation throughout.

We define the *uniform spectrum* as the mutation spectrum that accesses all possible mutations with equal probability. While not expected empirically, this spectrum is important theoretically as no mutational class is either over- or undersampled (*7, 8, 27*). For a mutation rate spectrum with *N* mutational categories, each category has frequency 1*/N* in the uniform mutation rate spectrum. On the other hand, the uniform absolute count spectrum depends on the genome content. Again, if mutational category *i* has *n_i_* opportunities to occur in the genome, the uniform absolute count spectrum is simply given by *n_i_/*Σ*n_i_*.

### Bias Reversal Measure

Consider a set of somatic mutations isolated from some cancer of interest. We seek to compare the spectrum of these mutations with the germline spectrum. In particular, we would like to quantify the extent to which the cancer spectrum reverses (rather than reinforces) the mutational bias observed in the germline. As previously demonstrated (*7,8*), the bias is reversed if a particular mutation frequency moves toward (or in fact past) the uniform frequency. Thus for example the unbiased absolute transition frequency is 0.33. If the transition frequency in the germline is 0.7, a transition frequency of either 0.6 or 0.2 are both in the direction that reverses the germline bias, whereas a change to 0.8 reinforces the germline bias.

Mutations that occur more frequently, in absolute terms, will have a greater effect on the degree to which the mutation bias is reversed. We therefore use absolute count spectra to compute the bias reversal. If we consider *n* mutational categories, the bias may be closer to the uniform level than the germline bias for some categories, but may be further away in others. Let *d_i_* represent, for the *i*th mutational category, the direction which constitutes a bias reversal, relative to the germline. Thus *d_i_* = 1 if the germline has a lower frequency than the uniform level, such that an increase in frequency represents a bias reversal. Similarly, *d_i_* = −1 when the germline has a higher frequency than the uniform level and reducing this frequency would yield a bias reversal. (In the unlikely case that the germline and the uniform levels are exactly equal, there is no bias to be reversed and therefore we take *d_i_* = 0.) The overall bias reversal of a given spectrum is then measured by the quantity

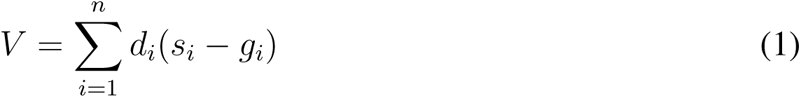

where *g_i_* and *s_i_* are the frequencies of the *i*th mutational category in the germline spectrum and the new spectrum, respectively.

Fig. S4 illustrates how the bias reversal measure is calculated for two fictional spectra with 3 types of mutations. We note two things: first, a spectrum need not reverse the bias in each individual mutational class to have a positive bias reversal measure; these effects are summed across classes. The extreme examples shown in this figure are for illustration. Second, we note that the bias reversal measure does not always combine linearly across scales. For example, the bias reversal measure at the Ts:Tv level will only be equal to the bias reversal measure at the 1-mer level if the frequencies of the two 1-mer Ti happen to shift, in every case, in the same direction as the overall Ti, and similarly for the four 1-mer Tv. When this occurs, the bias reversal measure combines linearly and is identical across scales, as observed in our data for example in fig. 1F.

### Signature Decomposition

Different mutational processes generate distinct combinations of mutation types. Recent efforts have successfully identified particular spectra, “mutational signatures”, associated with distinct mutational processes in cancer, and established links to their underlying mechanisms (*10*). A mutational spectrum from a particular tissue or tumor type can thus be decomposed into weighted contributions from known mutational signatures (*28–31*).

We use SigNet (*31*) to find the combination of mutational signatures, as identified in COSMIC v3.1 (*10*), that best describe the observed mutation counts in our data. This algorithm leverages the strong correlations between mutational processes observed in cancer data, providing highly accurate decompositions even when the number of mutations is low (*31*).

In brief: we first count the number of observed mutations for each of the 96 mutation types. Since COSMIC signatures are derived from whole-genome data, when decomposing mutational spectra from coding regions, SigNet first corrects for the 3-mer abundances in the coding genome; the input data is rescaled by the ratio between the 3-mer abundances in the coding genome and the whole genome.

The output of SigNet includes estimates of the signature weights, as well as a classification score, which reflects the degree to which the decomposition is considered reliable. SigNet also assigns a weight to an “unknown” category, which pools the weights from any signatures with predicted weight lower than 0.01.

### Positive Selection Detection in Cancer Genes

The ratio of non-synonymous to synonymous mutations (dN/dS) is a standard measure used to detect the influence of natural selection; positive selection is inferred when dN/dS*>*1, and genes with strong evidence for positive selection in tumour samples are considered cancer genes (*24*). Here, rather than detecting cancer genes, we use a similar approach to check whether HM and NHM samples show signs of positive selection in known cancer genes. In other words, we check if selection acts differently in HM and NHM using the pooled coding 3-mer spectra across tissues in each of these two categories. To avoid the dominance of skin and colon (see fig. S1A) in the pooled HM spectrum, mutations from skin and colon are downsampled so that the numbers of mutations from these two tissues are equal to the number of hypermutator mutations in the next most prevalent tissue, in this case, stomach cancers (fig. S1B).

Finding dN/dS is not possible for the four 3-mer mutation types (ACT*>*A, ACT*>*G, ATT*>*A and ATT*>*G) that do not generate synonymous mutations. Also, the number of synonymous mutations in cancer genes in our dataset is low for some tissue types (less than 200 mutations for 10 tissues out of 20). These data are clearly not sufficient to compose a 96-category 3-mer mutation spectrum.

Therefore, instead of using synonymous mutations, dS, as a neutral proxy, we use the fact that in the absence of positive selection, both cancer and passenger genes in tumour samples should have the same genomic mutation rates. This method allows us to calculate expectations for synonymous and non-synonymous mutations in all 3-mer contexts based on passenger genes, and compare these with the observed mutations in cancer genes. We interpret any observed differences as signs of positive selection because signals of negative selection are nearly absent in tumor evolution (but see (*25*) and (*32*)).

Consider a particular 3-mer. We first count the number of times this 3-mer occurs in the passenger genes in the human genome, *n_p_*. We then consider one of the three possible mutations of that 3-mer, and count the number of observed mutations of this type, in passenger genes, in the PCAWG dataset, *m_p_*. Dividing the observed mutation count in passenger genes by the number of occurrences of the 3-mer in passenger genes yields the expected genomic mutation rate for this 3-mer mutation, *µ* = *m_p_/n_p_*.

To determine the expected number of mutations of this 3-mer mutation type in cancer genes, we count the number of times this 3-mer occurs in cancer genes in the human genome, *n_c_*. In addition, we classify each of these instances as either synonymous or non-synonymous, depending on the outcome if this particular 3-mer mutation occurred at that position, such that *n_c_* = *n_cs_* + *n_cn_*, where *n_cs_* and *n_cn_* are the synonymous and non-synonymous instances respectively.

Given these values, it is straightforward to compute the expected numbers of synonymous and non-synonymous mutations in cancer genes, *e_cs_* = *µn_cs_* and *e_cn_* = *µn_cn_*, respectively. Finally, we count the number of observed mutations of this type in the PCAWG dataset in cancer genes, again differentiating synonymous (*m_cs_*) and non-synonymous (*m_cn_*) mutations such that *m_c_* = *m_cs_* + *m_cn_*. The relative difference between the observed and expected numbers of mutations is then a measure of whether there is an excess of a particular type of mutation in cancer genes. We refer to this relative difference as the “excess measure”, computed for each 3-mer mutation as:

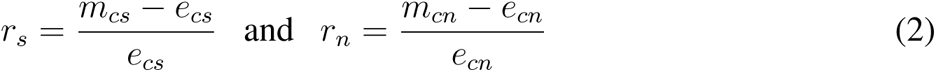

for synonymous and non-synonymous mutations, respectively. If the excess measure is positive for a given 3-mer mutation type, then more mutations of that type are sampled in cancer genes than in passenger genes, for example a value of 1 means there are twice as many mutations observed as expected. Due to the lack of synonymous mutations for the four 3-mer mutations mentioned above, the excess measures for synonymous mutations are defined for only 92 of the 96 mutation types.

We use bootstrapping to assess the statistical significance of the excess measure for all 3-mer mutation types in particular types of cancer. Since mutations in cancer genes form about 5% of all observed coding mutations in PCAWG, we bootstrap 50, 000 samples from all coding mutations, where each sample is of the same size (5% of the total pool of coding mutations). Each of these samples has a number of synonymous and non-synonymous mutations in each 3-mer context, forming the distributions *B_s_* and *B_n_* across the bootstrapped samples, respectively. We can then assign a *z*-score to the numbers of synonymous and non-synonymous mutations observed in cancer genes (*m_cs_* and *m_cn_*) assuming that the distributions *B_s_* and *B_n_* are approximately normal. Given that there are 96 comparisons, we use a conservative Bonferroni correction and consider an excess measure to be statistically significant at a *p*-value of 0.05*/*96 = 0.00052, which corresponds to the critical *z*-score value of 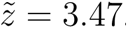.

## Results

### Reduced transition bias in non-hypermutated samples

We define the mutation bias of any mutational type to be reversed, compared to the germline bias, if it differs from the germline bias in the direction of the unbiased state, which assumes a uniform probability of each *k*-mer mutation type (see Methods) (*8*) (Fig. 1C). Using family-based datasets of *de novo* mutations (*22*), we find that the germline oversamples transitions (Ti) with a relative mutation rate (RMR) of 0.853 in coding mutations (Fig. 1D).

The Ti RMR in 9 normal tissue samples (Fig. 1D) is not significantly different from the germline (t-test, *p* = 0.074). The liver is the only outlier, with a Ti RMR of 0.55, presumably related to the transversion-rich mutational process induced by aristolochic acid exposure (SBS22 in COSMIC database), as observed in female donors (*19*). Similarly, the Ti RMR in the pooled data from HM cancers does not differ significantly from the Ti RMR in the germline (t-test, *p*=0.57). In contrast, NHM samples show a significantly reduced Ti RMR (t-test, *p* = 9.1 × 10^−9^).

These results are supported when the overall Ti RMR is decomposed at the 1-mer level (Fig. 1E). Note that the 1-mer RMRs of the germline are higher than the uniform spectrum level (1/6) for transitions and lower for transversions. Even when the germline RMR is very close to the uniform level (e.g. C*>*A mutations), non-hypermutators show strong evidence for a reversed bias; for both transitions they have a significantly reduced RMR compared to the germline, while two of four transversions show a significantly elevated RMR (stars; Bonferroni-corrrected *p <* 0.05*/*18 = 0.0028). In contrast, the hypermutator 1-mer mutation spectra differ significantly from the germline only for C*>*G transversions, where the Tv bias is not reduced but is in fact significantly reinforced. Note that none of the 1-mer mutation distributions for normal tissues have significantly different means from the germline. Analogous results hold for whole-genome mutations (Fig. S5).

To extend our analysis to the 3-mer level, we define a new metric, the bias reversal measure, to summarize spectrum changes across mutation types. For any spectrum, the bias reversal measure sums the degree to which the spectrum reverses the bias observed in the germline (see Methods). For passenger-gene (non-cancer-gene) mutations in HM and NHM cancers and at the Ti:Tv, 1-mer, and 3-mer levels, non-hypermutators show distributions of this measure across tissues that are significantly higher than zero (Fig. 1F; t-test, *p <* 2.8 × 10^−9^ in all three cases), whereas normal tissues and hypermutators are not significantly different from zero (t-test, *p >* 0.065 in all six cases), meaning that the bias is significantly reversed in NHM, but no such pattern is observed in normal tissues or HM. Similar results hold for the whole genome (Fig. S6).

We restricted our analyses of coding spectra to passenger genes to reduce possible confounding effects of positive selection (see Methods). Returning to cancer genes, we find that in normal tissue, HM, or NHM cancers, mutations in cancer driver genes tend to have a higher bias reversal than mutations in passenger genes (’x’ in Fig. 1F). This higher bias reversal for cancer genes relative to passenger genes is not driven by differences in the expected counts of 3-mer mutations, as these are highly correlated between these gene classes (*R >* 0.99).

### Non-hypermutator 3-mer spectrum is highly similar across tissues

The 3-mer mutation rate spectra for the HM and NHM samples, summed across tissues, are significantly positively correlated (Fig. 2A, Spearman rank correlation *R* = 0.54, *p*=1.89×10^−08^; also see Fig. S7 and Fig. S8 for the full 3-mer spectra). To further test for similarity among spectra, we computed the rank correlation coefficient between each tissue spectrum and all other spectra (pooled) in the same class. For instance, we correlate the spectrum from each tissue’s NHM samples with the spectrum computed by pooling NHM samples from all other tissues. NHM spectra are significantly more strongly correlated with one another (mean *R* = 0.86 ± sd = 0.052), than HM spectra are with one another (mean *R* = 0.62 ± sd = 0.16; Fig. 2B t-test, *p* = 7.8 × 10^−4^). Considering unpooled pairs of tissues, the correlation between spectra of NHM cancers is higher than HM cancers (Fig. 2C).

**Fig. 2:**
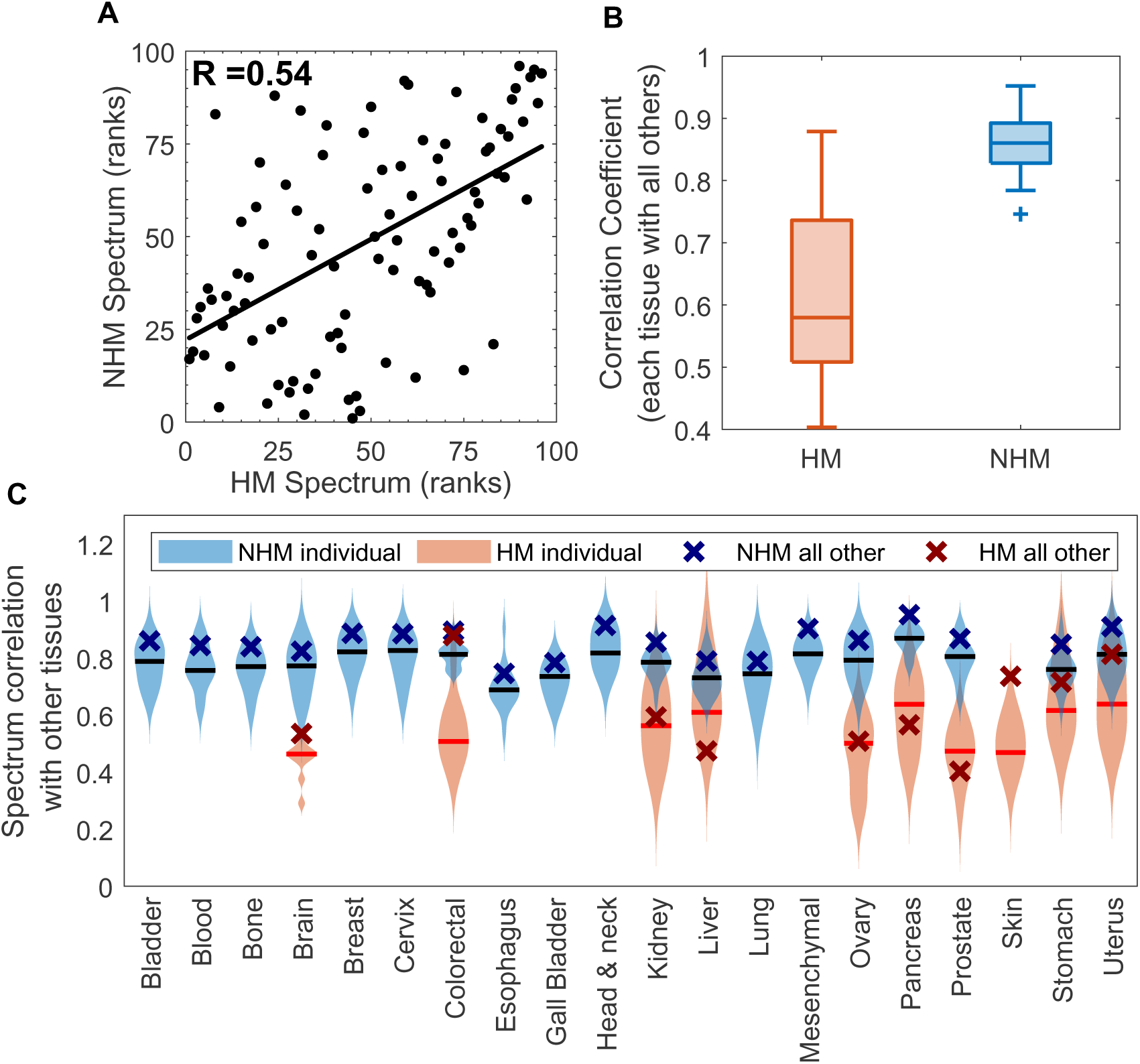
Non-hypermutator spectra are highly similar across tissues, unlike hypermutator spectra. (A) Correlation of the 3-mer spectra from NHM samples and HM samples; the Spearman correlation coefficient is *R* = 0.54. (B) Correlating the spectrum of each tissue with the pooled spectrum from all other tissues in the same class (HM or NHM) shows high Spearman correlation coefficients for NHM (blue) and lower values for HM (red). Centre bar indicates distribution median and whiskers show 95% CI; ’+’ signs represent outliers. (C) Spearman correlation coefficients of the spectrum of each tissue with each other tissue in the same class (violin plots); horizontal lines represent the medians of the distributions. Correlation coefficients between each tissue and pooled samples from all other tissues in the same class (’x’ symbols) are also shown for comparison; these are the values summarized in panel (B).

### What mutational processes characterize non-hypermutator tumors?

To examine the mutational processes underlying mutation spectra changes, we conducted a signature decomposition analysis (*31*)(Fig. 3A). For the pooled NHM spectrum, 52% of the mutations are attributed to mutation signature SBS5, 16.8% are related to the AID/APOBEC family of cytidine deaminases (SBS2 and SBS13), and 14.4% of the mutations are attributed to SBS40, while other signatures have lower weights. All of the signatures composing the NHM spectrum have positive bias reversal measures (Fig. 3B), excluding SBS1 and SBS2.

**Fig. 3:**
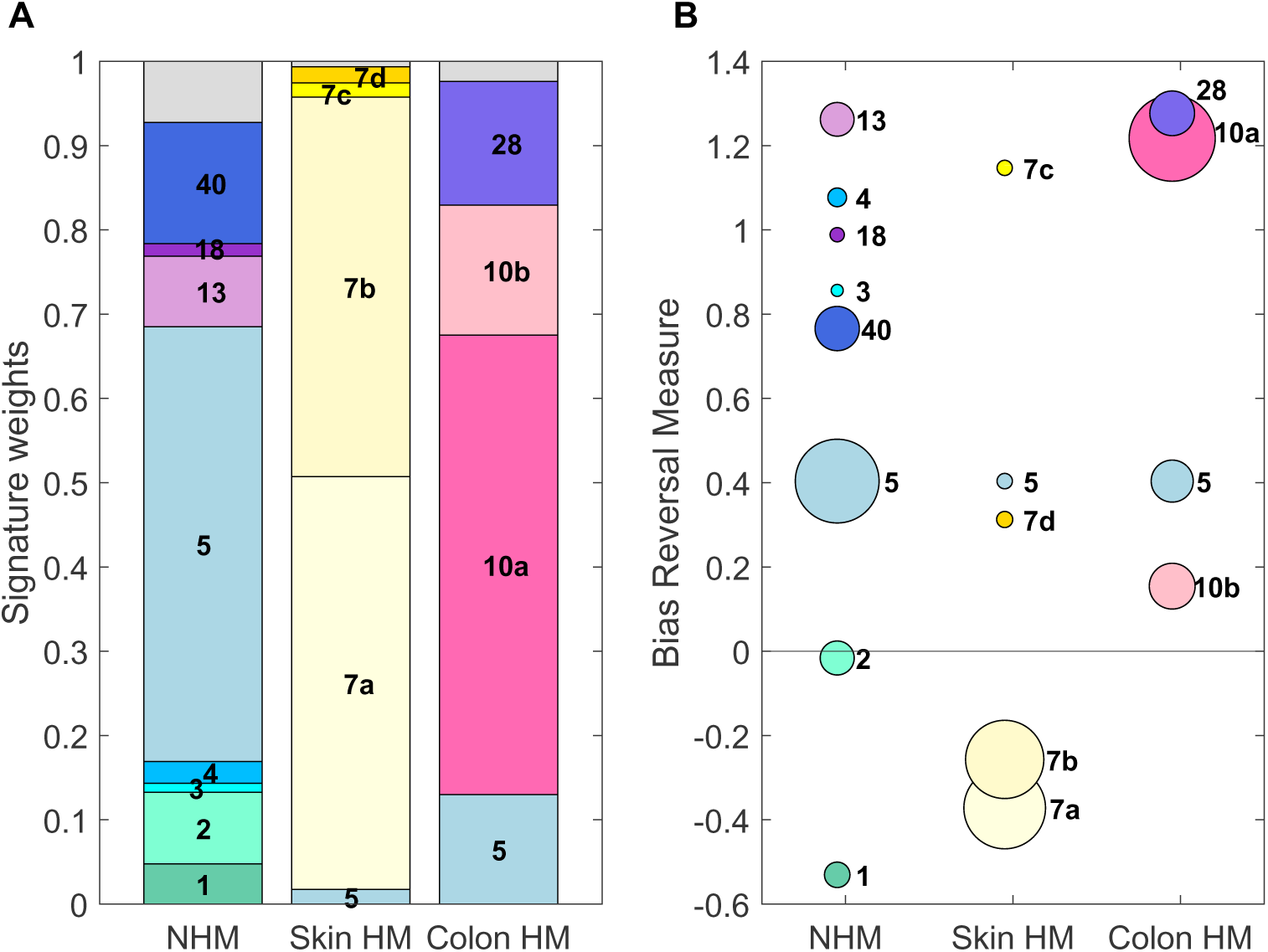
Most signatures composing the NHM and colon HM spectra reverse the bias, while mutational processes tend to reinforce the bias in skin HM samples. (A) The weights for each mutational signature identified as contributing to the three spectra are shown; grey proportions at the top represent mutations from unknown mutational processes. (B) Bias reversal measures for each signature; circles have an area proportional to the weight of that signature within the respective spectrum.

Since the spectra of HM tumors vary substantially across tissues (Fig. 2C), we aimed to perform tissue-specific signature decompositions of HM samples, but only skin and colon had sufficient mutations (Fig. S1, also see Fig. S8 for the full 3-mer spectra). As expected, UV light signatures (SBS7a through SBS7d) dominate the skin spectrum (Fig. 3A); the two dominant signatures, SBS7a and SBS7b, have a reinforced bias (negative bias reversal measure; Fig. 3B). In contrast, mutations derived from a defective polymerase epsilon dominate the colon HM spectrum (SBS10a and SBS10b, and the associated SBS28; Fig. 3A). All signatures underlying the colon HM spectrum have a positive bias reversal measure (Fig. 3B), explaining why colon cancer was the outlier with strong transition bias reversal in HM cancers (Fig. 1D).

### Positive selection in cancer genes in NHM cancers anticorrelated with the germline spectrum

Previous work has identified stronger signals of positive selection in tumors with lower mutation rates (*24, 32*). Our results suggest that mutations that are positively selected in cancer, at least in NHM, may reverse germline mutation biases. To examine positive selection in both HM and NHM tumors, we compute the excess measure (eq. 2) for each 3-mer mutation

(Fig. 4A. In both HM and NHM categories, we observe positive and negative excess measures for different 3-mer mutation types, but non-synonymous mutations in NHM show the highest number of positive values, with a distribution mean that is significantly different from zero (Fig. 4A; t-test, *p* = 2.02 × 10^−21^). The non-synonymous distribution in NHM is significantly different from each of the other three distributions (Wilcoxon rank sum test, *p <* 3.04 × 10^−5^). In agreement with previous work (*24, 32*), we thus find stronger evidence for positive selection acting on non-synonymous mutations in NHM than in HM tumors. Note that the distribution mean for non-synonymous mutations in HM is also significantly different from zero (t-test, *p* = 0.0065), but the distribution means for synonymous mutations are not significantly different from zero (t-test, Bonferroni-corrected, *p* = 0.44 and *p* = 0.04 respectively).

**Fig. 4:**
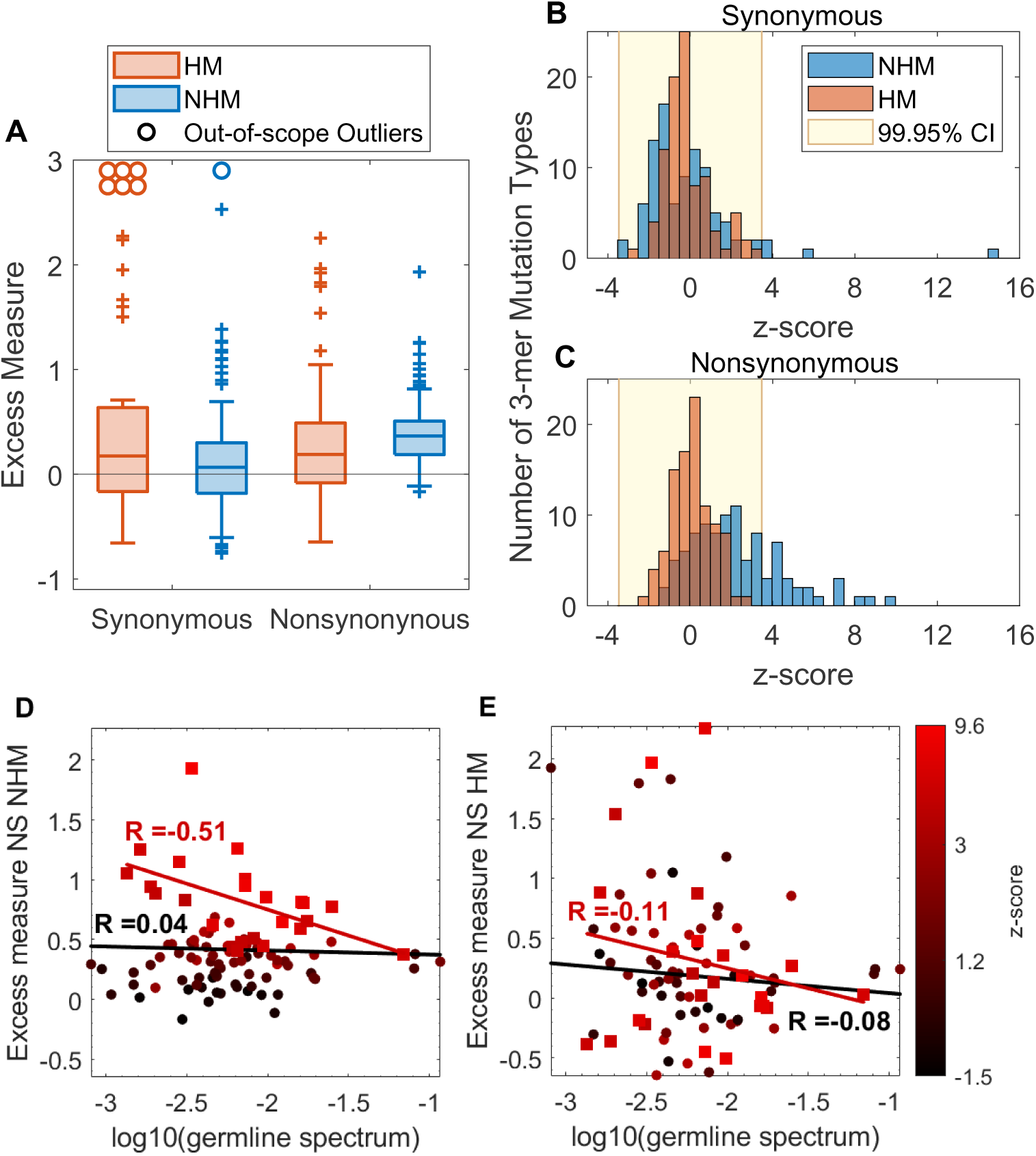
Evidence of positive selection in cancer genes is stronger in non-hypermutators than hypermutators. (A) Excess measures comparing cancer and non-cancer genes for all 3-mer mutation types are shown for synonymous and non-synonymous mutations in both HM (red) and NHM (blue). Only the non-synonymous distributions are significantly higher than zero. Boxplot whiskers include 95% CI. The positive *y*-axis is truncated for clarity; ’+’ signs represent outliers. (B) Histograms of z-scores obtained by comparing the numbers of synonymous mutations of each 3-mer mutation type in cancer genes with results in 50,000 bootstrapped samples from all synonymous mutations. The Bonferroni-corrected CI from the boostrapped dataset is shown in yellow. (C) Analogous results for non-synonymous mutations. (D,E) Excess measure for non-synonymous mutations in NHM and HM is correlated with the germline non-synonymous passenger-gene spectrum (log-scale). The 3-mer mutations that show significant positive selection for NS NHM (out of CI in panel (C), squares) show a significant anti-correlation only in the NHM case (red lines, (D):*p*=0.012, (E):*p*=0.61), while all 3-mer mutations taken together (squares and circles) do not show any significant correlations (black lines, (D):*p*=0.7, (E):*p*=0.43). The color of ea^2^c^1^h point corresponds to the NHM z-score from panel (C). Open circles/squares (in panels A and E) indicate out-of-scope outliers.

To detect selection on individual 3-mer mutation types (and to account for the variance in excess measure for rare mutation types), we compared the numbers of mutations observed in cancer genes with bootstrapped samples from all coding mutations. For all 3-mer contexts in HM, and almost all those in NHM, the number of synonymous mutations in cancer genes falls within the 99.95% confidence interval of the bootstrapped samples, showing no sign of selection (Fig. 4B). In contrast, non-synonymous mutations show 24 positively-selected 3-mer mutation types in NHM (out of 96), but none in HM (fig. 4C). Non-synonymous excess measures in these 24 3-mer mutation types in NHM (Fig. 4D) are anti-correlated with the corresponding mutation count frequencies in the non-synonymous germline spectrum, while this is not true for HM cancers (Fig. 4E), demonstrating that among mutations under positive selection, those least likely to occur in the germline show the strongest selective effect in NHM. There is no significant correlation, however, between the germline mutation spectrum and the measure of positive selection across all 3-mer mutation types.

Negative selection is typically detectable only in tumors with a very low mutation burden, as expected under strong clonal interference (*32*); consistent with this expectation, we found no 3-mer mutation type with an observed value that falls below the confidence interval, in HM or in NHM (Fig. 4B).

## Discussion

When looking at cancer through an evolutionary lens, cancer-driver mutations are “beneficial” to the cell, as they increase a cell’s reproductive rate, leading to local or distant invasions (*1,33*). These driver mutations are prevalent across cancer types, but rare in normal tissues (*34*). While some cancers may develop a mutator phenotype, which can explain their increased access to driver mutations, a mutator phenotype is not essential for carcinogenesis (*35*). Our previous work (*8*) suggests another mechanism that enhances access to previously rare mutations, that is mutational bias shifts. Here we tested whether such bias shifts exist in cancer, and if so, whether this occurs only in non-hypermutated cancers or also in hypermutators.

We examined the mutational spectra of thousands of human tumor samples, computing spectra for hypermutator (HM) and non-hypermutator (NHM) cancers across twenty tissue types. We demonstrate that the mutational spectra in NHM tumors is highly correlated across tissues, an effect not observed in HM tumors (Fig. 2). Thus while HM tumors show a dramatic increase in mutation rate, NHM tumors show only modest increases in mutation rate but have a distinct mutation spectrum that is repeated across diverse tissues and donors.

Reversals of mutation bias, that push the mutation spectrum either towards or past the unbiased/uniform state, offer access to undersampled classes of mutations (*7, 8, 27*), which we hypothesized may include the mutations that drive cancer. Consistent with this prediction, whether analyzed at the Ts:Tv, 1-mer or 3-mer levels, the distinct mutation spectrum of NHM (Fig. 2) tumors significantly reverses the germline bias (Fig. 1), while HM tumors and normal tissues show no such effect. These results suggest that while HM tumors access cancer driver mutations through elevated mutation rate, NHM tumors access driver mutations through changes in mutation spectrum that correspond to reversals in germline mutational biases (Fig. 5).

**Fig. 5:**
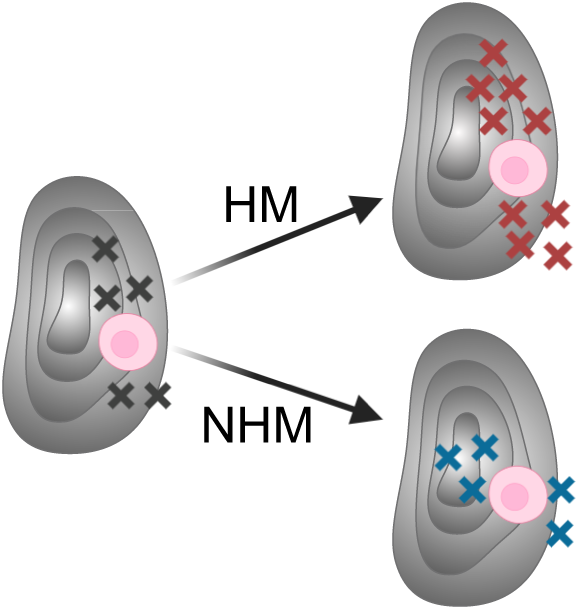
Cancer driver mutations are rarely accessed but “beneficial” in the fitness landscape (grey surface) of a cellular lineage. These can be accessed by two possible changes in mutational processes: an increase in mutation rates as seen in hypermutators (HM), or a reversal in the mutational bias as seen in non-hypermutators (NHM).

We further confirm this hypothesis by demonstrating that driver mutations in NHM are indeed stronger, and/or more common, for those types of mutations that are less likely to occur in the germline or in healthy tissues (Fig. 4D). This suggests that indirect selection on 3-mer mutation modifiers (*36*) may tune the human germline and healthy somatic mutation spectrum such that the greater the effect of a cancer driver mutation, the less likely it is to occur, and if it does occur, the less likely it is to remain unrepaired.

Our signature decomposition analysis (Fig. 3) suggests SBS13 and SBS40 as mutational processes that reverse the germline bias and play a key role in the NHM mutation spectrum. While SBS5 makes the largest contribution to the NHM spectrum, it is a clock-like signature (prevalence increases with patient age) that shows a modest bias reversal and is found in the germline and healthy tissues as well. In contrast, SBS13 is attributed to activity of the AID/APOBEC family of cytidine deaminases (*37*), which is known to promote carcinogenesis and cause genomic instability (*38–40*). This mutational signature has been previously detected in many cancer types (see Fig. 3 in (*10*)), and it reverses the germline bias mainly due to its richness in C*>*G transversions. Likewise, SBS40 is a mutational signature detectable in many cancers (*10*), and it strongly reverses the germline transition bias because it mimics a uniform spectrum (Fig. 3). There is a significant positive correlation between SBS40 and hypoxia (*31*), and the presence of transient or chronic hypoxia, characterized by critically low tissue oxygen, is a defining characteristic of cancer (*41*). This hypoxic state is associated with a significant increase in mutation rate, thought to result from a reduction in the effectiveness of various DNA repair mechanisms (*42, 43*).

Further work needs to be done to test potential temporal contributions of these two mutational signatures to tumorogenesis. While the activity of the AID/APOBEC family is mutagenic, likely contributing to initial tumor formation (*38, 39*), many APOBEC3-induced mutations occur later in tumor evolution (*44*). Similarly, hypoxic conditions in the healthy tissue, due to inflammation, could also precede tumourigenesis (*45*). Once a solid tumor has developed a certain minimum size, we hypothesize that the hypoxic state could enable a second round of driver mutations that are required for metabolic changes and angiogenesis allowing the tumor to survive and grow under these new conditions. Further studies are also needed to determine whether the distinct and remarkably stable mutation spectrum observed in NHM tumors is due to alterations in DNA repair enzymes resulting from common hypoxic conditions among patients, from other unknown causes shared among patients, or from multifactorial convergent evolution at the mutation spectrum level.

In contrast with the NHM spectrum, we observed a wide variability in the spectra of HM tumors. While skin cancers show no evidence of reversing the germline bias, HM samples from colon show a strong bias reversal. Simultaneously reversing mutation bias and increasing mutation rate provides more access to beneficial mutations than either alone (*8*). We speculate that the high incidence of colon cancer could be in part due to this powerful combination of an increased mutation rate along with a reversal of the germline bias. Interestingly, it has been observed that mutations in hypermutated and non-hypermutated colon cancers target different cancer genes (*46*), and here we show that colon cancer has two different spectra in these two categories.

By comparing observed and expected mutation counts, we show that NHM tumors have an excess of non-synonymous mutations in cancer genes in several 3-mer contexts, which we interpret as evidence for positive selection (*24, 25, 47*). This excess is significant not only for non-synonymous mutations, but for synonymous mutations in a few contexts, which is not completely unexpected since several synonymous mutations have been previously identified to act as cancer driver mutations due to their impact on splicing, RNA secondary structure and expression levels (*48, 49*).

On the other hand, we find no strong signal for positive selection in HM samples. This could be due to the combined effects of clonal interference (*32, 50*) and deleterious load in HM tumors, such that only a small fraction of mutations will successfully spread through the population. This effect could dilute signals of positive selection when mutation rates are very high. Since the small fraction of beneficial mutations that spread are also expected to have very large selective effects, negative epistasis could also reduce the signals of positive selection in HM tumors. This can occur when a single, large-effect mutation eliminates the need for further mutations that would otherwise be beneficial, for example when cancer driver genes “break” the same regulatory pathway. Moreover, HM tumors are known to have longer sequence context dependencies than those accounted for by our 3-mer model (*47, 51*). This may also make our detection of positive selection across 3-mer mutations less reliable in HM tumors than in NHM tumors. Nevertheless, we find that the excess of non-synonymous mutations in the 24 3-mer mutation types that show positive selection in NHM is also anticorrelated with the germline mutation spectrum in HM tumors (Fig. 4D,E). This suggests that the underlying fitness landscape may be quite similar for HM and NHM, but the inference of positive selection is simply more difficult in HM.

Our work putatively identified 24 3-mer mutation types that show positive selection in NHM. Classifying the contribution of these mutations to oncogenes and tumor suppressor genes is a clear avenue for future work, as is characterizing the distribution of fitness effects of these mutation types in the germline. More generally, we hope that characterizing the distinct 3-mer spectrum observed in NHM cancers may shed further light on cancer driver mutations and their critical role in early oncogenesis.

## Acknowledgments

**Funding:** This work was supported by the Natural Sciences and Engineering Research Council of Canada grant RGPIN-2019-06294, by the National Institute of General Medical Sciences of the National Institutes of Health through grants R01GM127348 and R35GM149235, and by the Spanish Ministry of Science and Innovation through the Centro de Excelencia Severo Ochoa (CEX2020-001049-S, MCIN/AEI /10.13039/501100011033), the Generalitat de Catalunya through the CERCA programme, and the European Union’s H2020 research and innovation programme under Marie Sklodowska-Curie grant agreement No.754422.

**Authors’ contributions:** MZT, DC, RNG and LMW designed the study. MZT and DC performed data analysis; CSC performed spectral decompositions. All authors contributed to data interpretation. MZT, LMW and RNG created the figures. MZT and DC drafted the manuscript. All authors edited and finalized the manuscript.

**Competing interests:** none.

## Data and materials availability

Functional annotations per coding site using SnpEff and Supplementary tables can be found in https://github.com/MarwaTuffaha/CancerBiases. Data are reported in the supplementary tables. Tables S1 to S4 provide data shown in figures 1 to 4 in the main text, respectively. Table S5 provides the 3-mer spectra plotted in figures S7 and S8.

## Supplementary materials

Figures S1 to S8

Tables S1 to S5

