## Supplementary figures for "Non-hypermutator cancers access driver mutations through reversals in germline mutational bias"


**The PDF file includes:**

Figs. S1 to S8

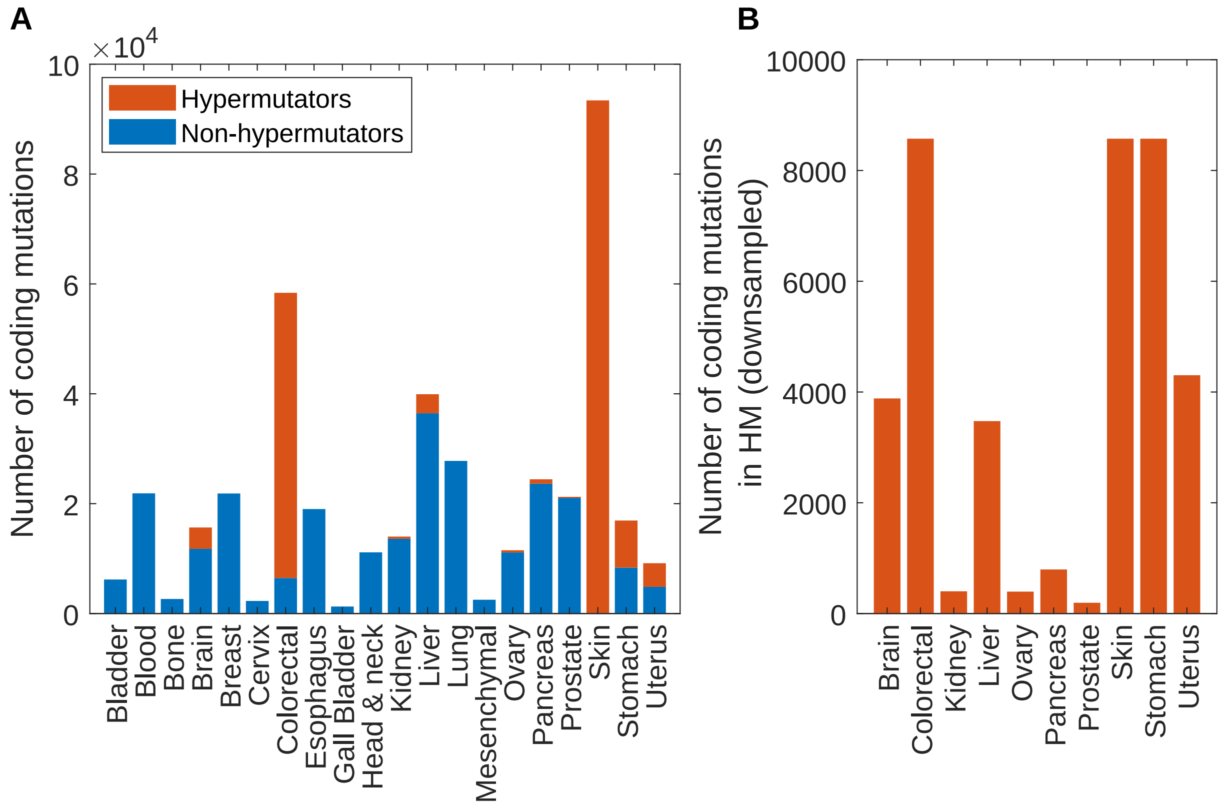


Figure S1: (A) The number of coding mutations in the PCAWG dataset, partitioned by tissue. Mutations from hypermutated and non-hypermutated samples are shown in red and blue, respectively. (B) The number of coding mutations in the hypermutated data set, where skin and colon mutations are downsampled such that the numbers of mutations from these two tissues are equal to the number of hypermutator mutations in the next most prevalent tissue (stomach).

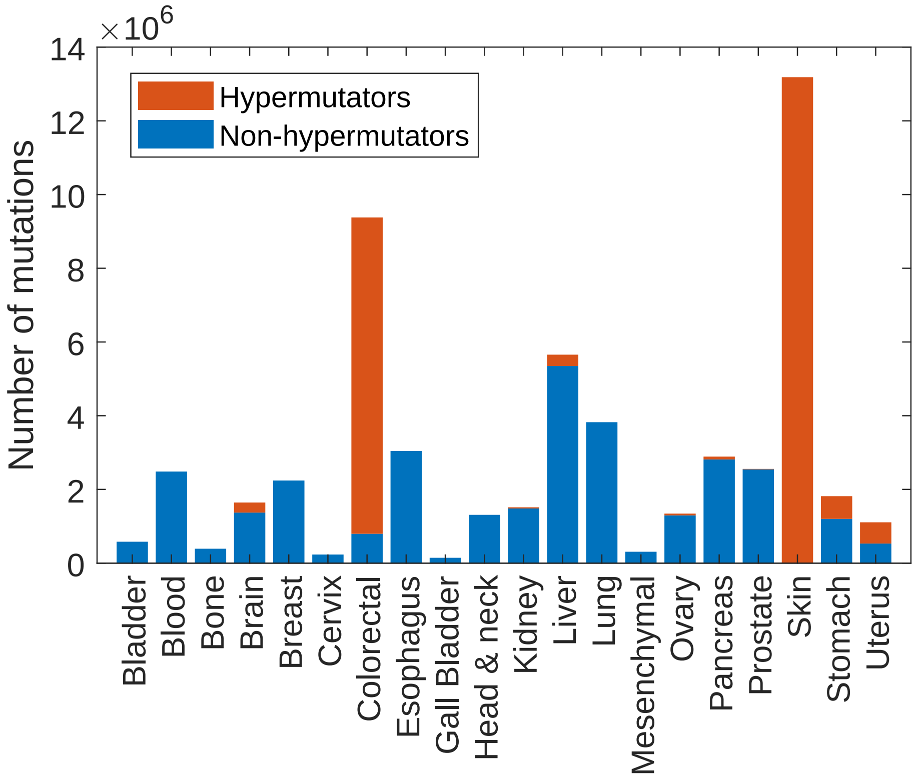


Figure S2: The number of whole-genome mutations in the PCAWG dataset, partitioned by tissue. Mutations from hypermutated and non-hypermutated samples are shown in red and blue, respectively.

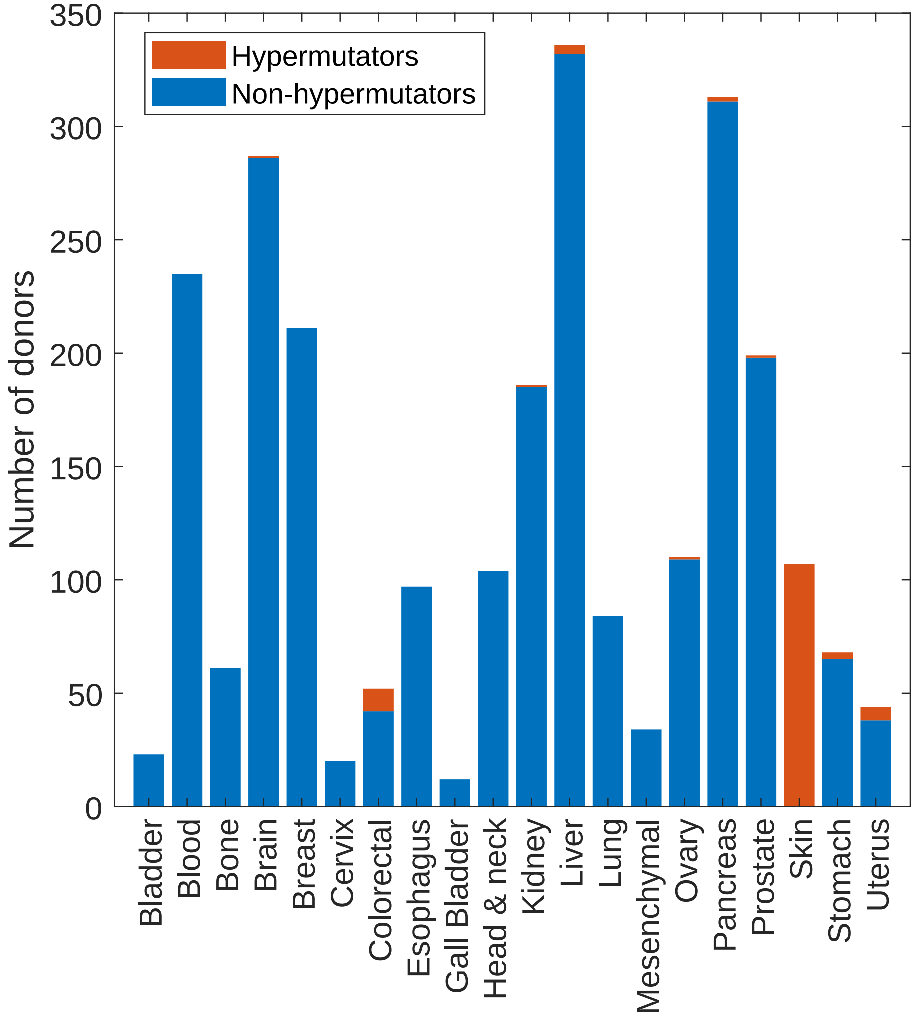


Figure S3: The number of donors (i.e. tissue samples) in the PCAWG dataset, partitioned by tissue. Hypermutated and non-hypermutated samples are shown in red and blue, respectively.

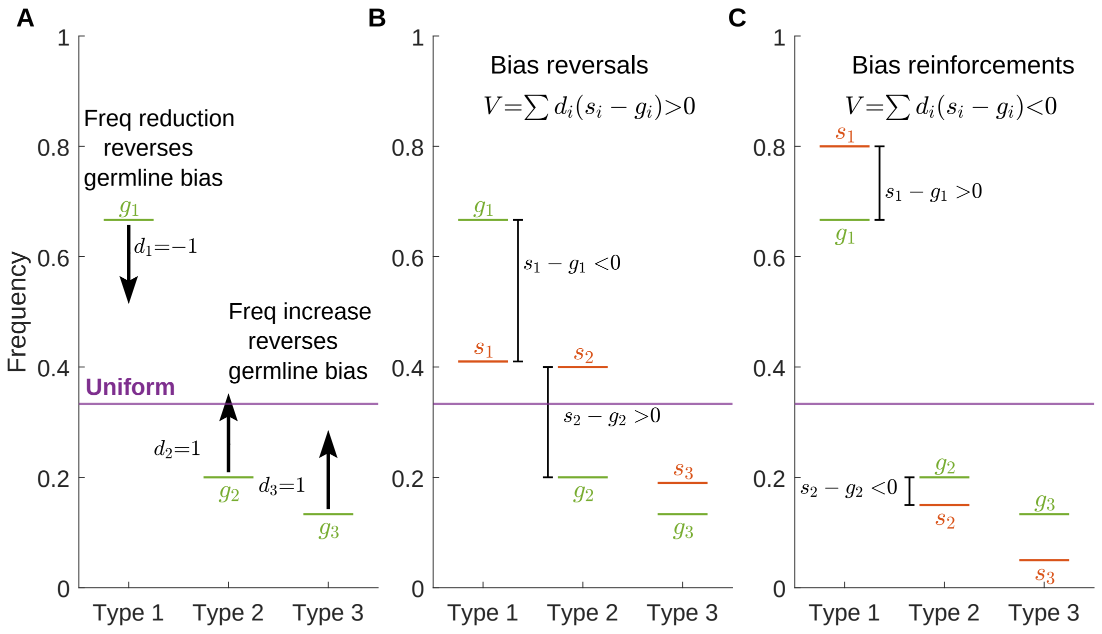


Figure S4: Example of fictional spectra with 3 types of mutations that all have the same occurrence opportunity (uniform frequency) in the genome of 1/3. Type 1 is over-represented in the germline ($g_1$, green) while the other two types are under-represented. (A) Arrows show the direction that reverses the bias (from germline frequency toward the uniform frequency) for each mutational type. (B) Example of a spectrum ($s_i$, red) that reverses the bias for all three mutation types, and thus has a positive bias reversal measure. (C) Example of a spectrum that reinforces the bias and thus has a negative bias reversal measure.

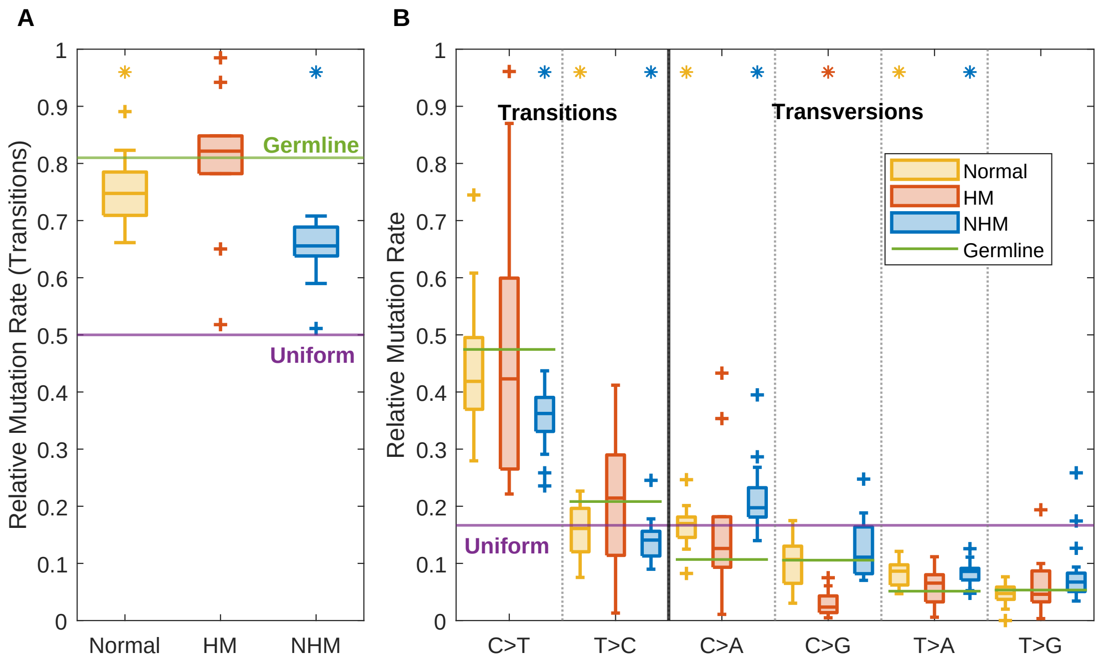


Figure S5: Results for whole genome spectra analysis are analogous to coding spectra analysis in fig. 1D,E. (A) Distributions of the RMR for transitions in normal tissues (yellow), HM (red) and NHM (blue) in different tissues, compared to the uniform and germline levels (purple and green lines, respectively). (B) The corresponding RMRs for each 1-mer mutation type. Arrows show the direction that reverses the bias for each mutational type. Horizontal bars within box-plots indicate medians; whiskers indicate the $95\%$ confidence interval; '+' symbols represent outliers. Stars indicate distribution means that are significantly different from the germline.

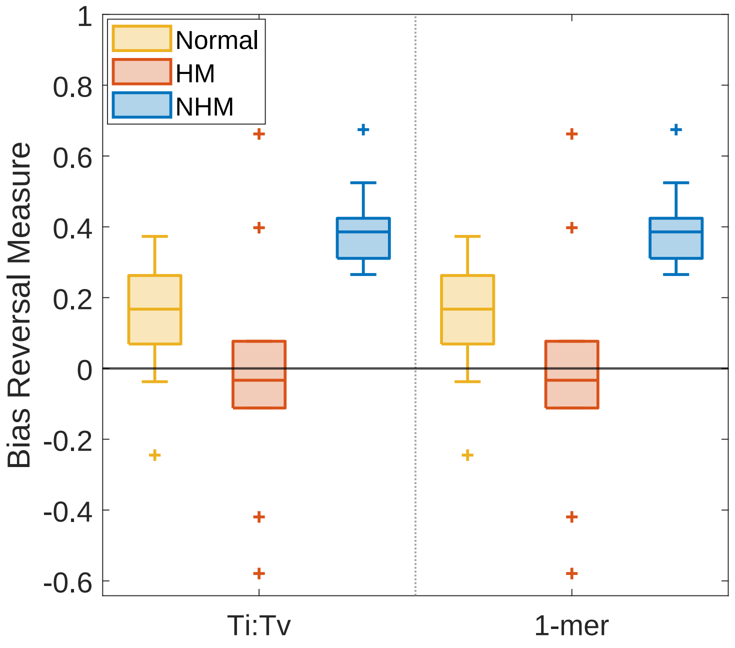


Figure S6: Bias reversals ($y$-axis) for whole-genome spectra in different tissues (boxplots) are significantly higher in NHM than in HM or normal tissues on the transition:transversion and 1-mer levels of analysis ($x$-axis). Horizontal bars within boxplots indicate medians; whiskers indicate the $95\%$ confidence interval; open circles represent outliers.

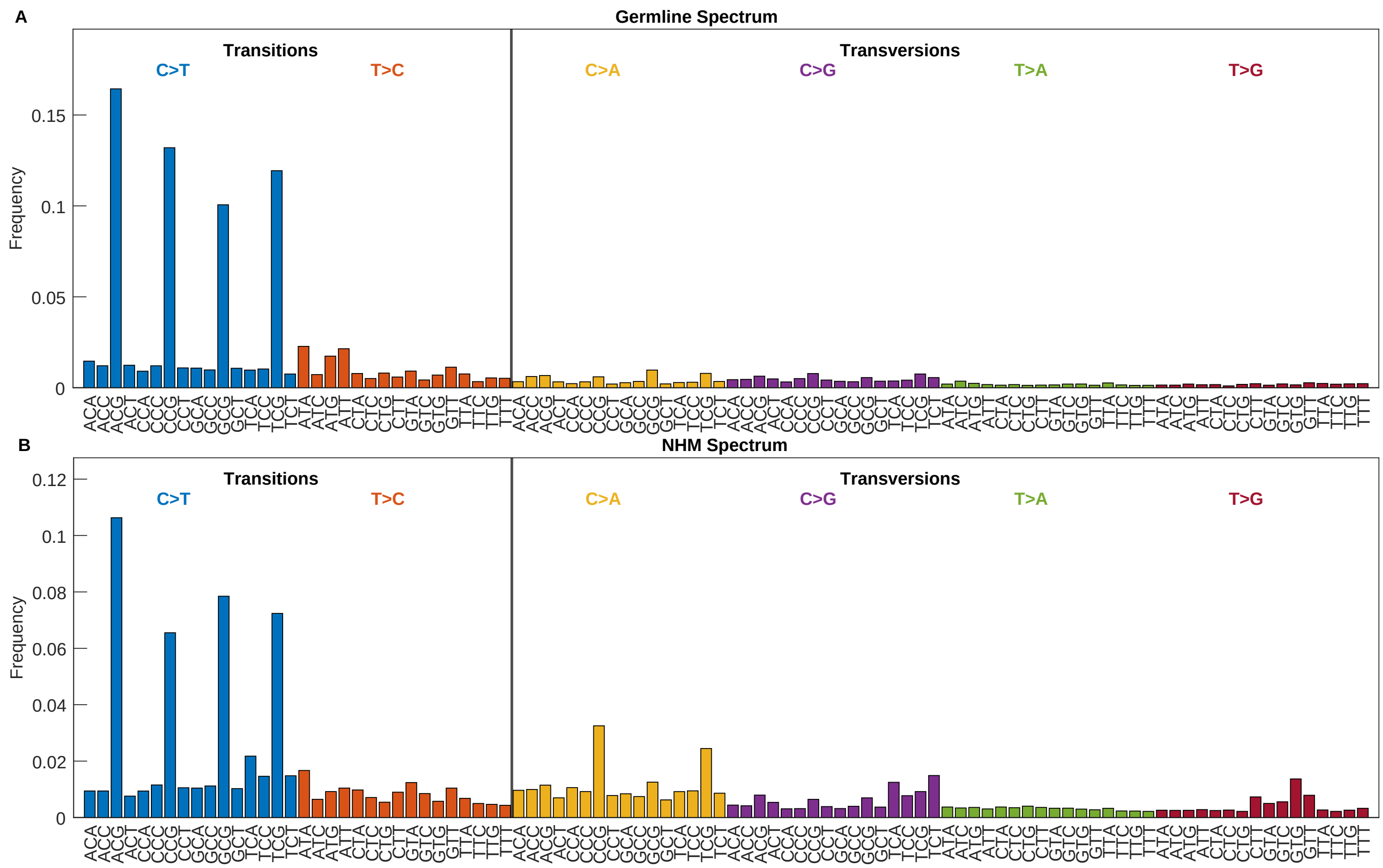


Figure S7: Mutation rate spectra in (A) germline and (B) non-hypermutators.

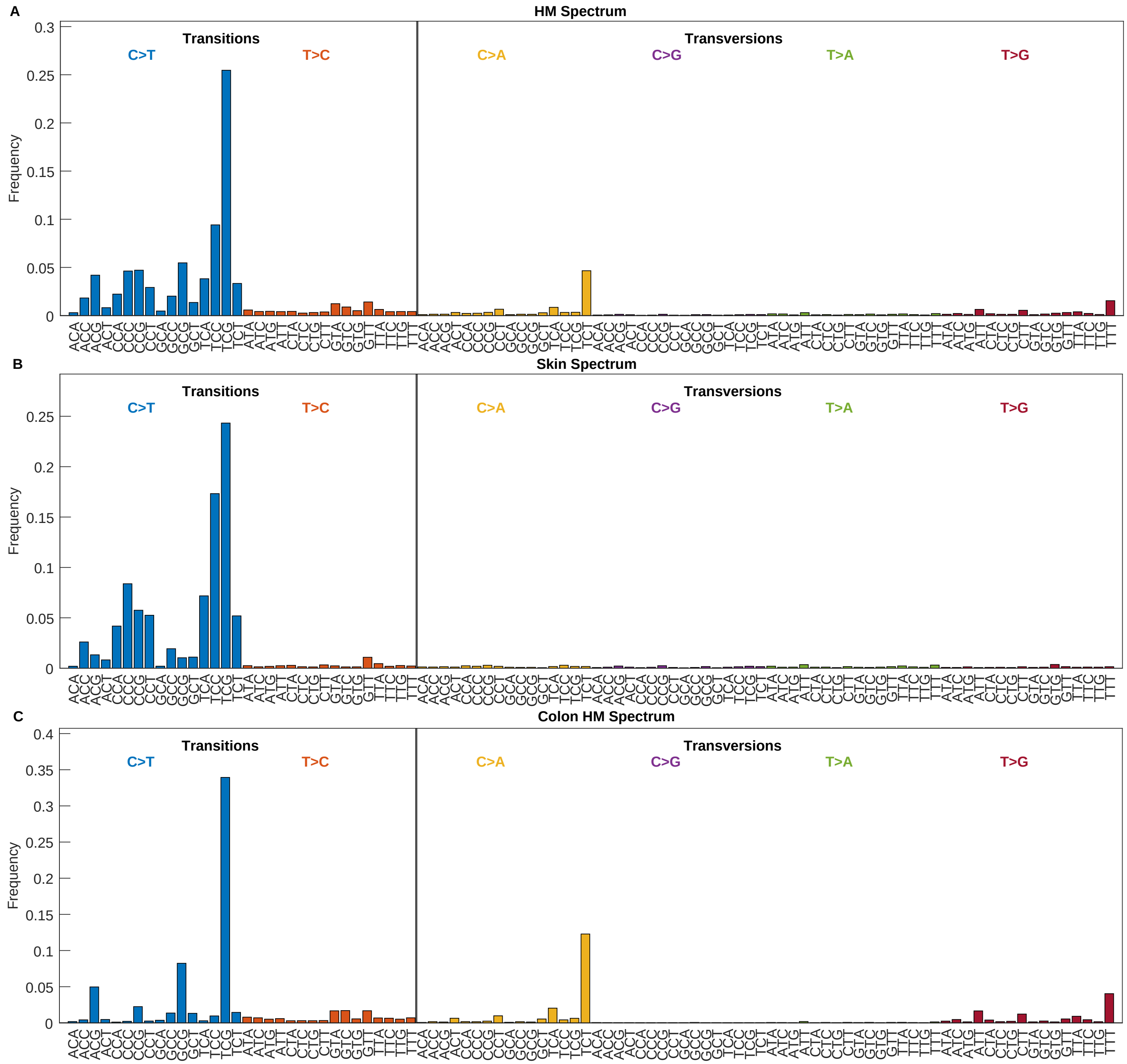


Figure S8: Mutation rate spectra in (A) pooled hyper-mutated samples, (B) skin samples (which are all HM), and (C) hypermutated samples from colon.
